# DNA methyltransferase enhanced *Fusobacterium nucleatum* genetics

**DOI:** 10.1101/2022.07.20.500824

**Authors:** Ariana Umaña, Tam T.D. Nguyen, Blake E. Sanders, Kevin J. Williams, Bryce Wozniak, Daniel J. Slade

**Affiliations:** Virginia Polytechnic Institute and State University, Department of Biochemistry, Blacksburg, VA, USA

**Keywords:** *Fusobacterium*, *Fusobacterium nucleatum*, DNA methyltransferase, methylase, methylome, restriction-modification, R-M systems, transformation, bacterial genetics

## Abstract

Bacterial restriction-modification (R-M) systems are a first line immune defense against foreign DNA from viruses and other bacteria. While R-M systems are critical in maintaining genome integrity, R-M nucleases unfortunately present significant barriers to targeted genetic modification. Bacteria of the genus *Fusobacterium* are oral, Gram-negative, anaerobic, opportunistic pathogens that are implicated in the progression and severity of multiple cancers and tissue infections, yet our understanding of their direct roles in disease have been severely hindered by their genetic recalcitrance. Here, we demonstrate a path to overcome these barriers in *Fusobacterium* by using native DNA methylation as a host mimicry strategy to bypass R-M system cleavage of user introduced plasmid DNA. We report the identification, characterization, and successful use of *Fusobacterium nucleatum* (*Fn*) Type II and III DNA methyltransferase (DMTase) enzymes to produce a multi-fold increase in gene knockout efficiency in the strain *Fusobacterium nucleatum* subsp. *nucleatum 23726* (*Fnn* 23726), as well as the first efficient gene knockouts and complementations in *Fnn* 25586. We show plasmid protection can be accomplished *in vitro* with purified enzymes, as well as *in vivo* in an *E. coli* host that constitutively expresses *Fnn* DMTase enzymes. By characterizing specific DMTases that are critical for bypassing R-M systems, we have enhanced our understanding of potential enzyme combinations, with the goal of expanding these studies to genetically modify clinical isolates of *Fusobacterium* that have thus far been inaccessible to molecular characterization. This proof-of-concept study provides a roadmap to guide molecular microbiology efforts of the scientific community to facilitate the discovery of new *Fusobacterium* virulence genes, thereby leading to a new era of characterizing how an oral opportunistic pathogen contributes to an array of human infections and diseases.

**IMPORTANCE:** *Fusobacterium nucleatum* is an oral opportunistic pathogen associated with diseases including cancer and preterm birth. Our understanding of how this bacterium modulates human disease has been hindered by a lack of genetic systems. Here we show that *F. nucleatum* DNA methyltransferase modified plasmid DNA overcomes the transformation barrier and allows the development of genetic systems in previously inaccessible strains. We present a strategy that can be expanded to enable the genetic modification of clinical isolates, thereby fostering investigational studies to uncover novel host-pathogen interactions in *Fusobacterium*.

## INTRODUCTION

Bacteria have multiple mechanisms to keep out foreign DNA elements including physical barriers in the form of membranes, and innate and adaptive nucleotide recognizing systems to degrade foreign DNA before costly genome integration^1-3^. This ability to recognize self-versus non-self DNA is critical for productive genetic exchanges through horizontal gene transfers (HGT) between close species to receive adaptive advantages ^4-6^. The two main nucleic acid surveillance systems bacteria deploy are restriction modification (R-M systems) and CRISPR-Cas (clustered, regularly interspaced palindromic repeat-CRISPR-associated proteins) systems. In addition, a new system known as DISARM has joined the bacterial arsenal of DNA defense systems^7^. CRISPR-Cas systems are considered adaptive immune components because of their ability to chromosomally integrate foreign (i.e., viral) DNA to create memory for subsequent encounters^8-11^. In addition, rather newly characterized BREX (Bacteriophage Exclusion) systems exists in 10% of the sequenced bacterial genomes and block phage DNA replication and lysogeny in infected cells^12,13^. BREX differentiates itself from R-M systems in that phage DNA is not cleaved or digested, which suggests a unique bacterial defense system. While R-M systems serve bacteria well in their survival and adaptation, they present significant challenges for researchers aiming to understand these organisms through genetic manipulation in the form of gene knockouts. This genetic recalcitrance is widespread throughout the bacterial kingdom, and in many cases, leads researchers to gravitate towards using strains that have robust genetic systems, instead of the strains they truly want to study which have strong R-M system barriers.

R-M systems consist of restriction endonucleases (REases) and DNA methyltransferases (DMTase), which can either exist as a paired REase/DMTase operon that can also contain additional specificity genes, or lone DMTase genes^14-16^. The system works when REases cleave DNA that does not have the proper DMTase induced methylation sequences, thereby signaling to the bacteria that the detected DNA is foreign and unwanted. R-M systems are classified as Type I, II, III or IV according to their molecular structure, subunit composition, cleavage position, restriction site, and cofactor specifications (**Fig S1)**. Type I (genes *hsdRMS*) cuts exogenous DNA by forming protein complexes and random cleavage usually happens at substantial distances from an asymmetric recognition sequence (400 to 7,000 bp)^17^, while Type II consists of an individual restriction endonuclease and methyltransferase that cleave DNA at symmetrical recognition sites^18^. In a similar way to Type I, Type III forms a protein complex necessary for the restriction enzyme activity; however, the methyltransferase can function independently. DNA cleavage for Type III RM systems takes place 25 to 27 bp 3′ to an asymmetrical recognition sequence that is 5 to 6 bp in length^19^.Furthermore, Type IV systems asymmetrical recognize DNA sequences, and cleavage by REases at a defined distance from the recognition siteres. In addition, some of these systems contain multiple DMTases that can be adenine or cytosine specific, as well as the REase oddly showing methyltransferase activity^17,20-22^.

*Fusobacterium*, especially the species *Fusobacterium nucleatum (Fn)*, has garnered significant attention since this bacterium was reported to be overrepresented in colorectal cancer tumors more than a decade ago^23-25^. Classical studies mainly focused on the role of *Fn* in oral infections and diseases including periodontitis^26,27^, severe organ infections^28-31^, and preterm birth^32-34^. The majority of recent studies have shifted to focus on a potential direct causal role in adverse cancer phenotypes including heightened inflammation^35-37^, production of a carcinogenic metabolite^38^, induced metastasis^39-41^, DNA damage^42-44^, increased resistance to frontline chemotherapy drugs^45,46^, and overall worse patient prognosis^35,47,48^. Despite an increasing interest in understanding how this bacterium contributes to cancer, there are very few mechanistic studies of specific bacterial effector genes due to R-M system induced genetic recalcitrance. Because of this, our current molecular studies have been limited to *a few Fusobacterium strains that are able to* acquire ‘naked’ DNA and incorporate it into their genome by recombination with homologous sequences or, in the case of episomal multi-copy plasmids, by establishing a new episome. Of these are *Fusobacterium nucleatum* subsp. *nucleatum* 23726 (*Fnn* 23726; transformation by electroporation)^49-51^, *Fusobacterium nucleatum* subsp. *polymorphum (Fnp* 10953; transformation by electroporation)^52^, *Fusobacterium nucleatum* subsp. *polymorphum* 12230 (*Fnp* 12230; transformation by sonoporation)^53^, and a recent paper highlighting the first gene interruption in *Fusobacterium necrophorum* using DNA conjugation from *E. coli*^54,55^. Needless to say, these four strains do not encompass all of the *Fusobacterium* subspecies and their respective infections and diseases that we would like to study and highlights the need for molecular biology and biochemical studies to achieve universal genetics.

Seminal studies have successfully used DMTases to modify and protect plasmid DNA to facilitate molecular genetics in several other bacteria^56-58^. What we currently know about the R-M systems of *Fusobacterium* largely exist as bioinformatic predictions based on DMTase classification in the REBASE database^59^. However, this bioinformatic classification in most cases does not come with experimental DNA methylation analyses to match enzymes with their target sequences. Additionally, even when a DMTase is matched with its recognition and methylation sequence, this does guarantee that these modifications will be important for effectively protecting and transforming plasmid DNA. Therefore, the goal of this study was to biochemically characterize and utilize a broad range of *Fn* DMTases in host-mimicry by methylation to accelerate bacterial genetics in previously inaccessible strains. This technique has been used successfully in many studies but was coined Plasmid Artificial Modification (PAM) where it was used to enhance transformation in *Bifidobacterium adolescentis*^*60*^. *We* successfully report the use of *Fn* DMTase enzymes produced in *E. coli* to protect plasmid DNA, facilitating a significant increase in chromosomal incorporation of plasmid and transposons in multiple *Fn* strains, as well as the development of the first gene deletions in *Fnn* 25586. Our study is not exhaustive because of the sheer number of strains and enzymes that could have been tested, but we believe our successful strategies will provide a flexible roadmap for the scientific community to adopt DMTase based methods for genetic manipulation in *Fusobacterium*.

## RESULTS

### Bioinformatic identification and classification of R-M systems in *Fusobacterium*

As shown in **Figure 1A**, bacterial R-M systems act by blocking exogenous DNA from entering and being incorporated into the genome by digesting foreign, improperly methylated DNA that does not contain the ‘password’ for safe entry. Scientists have exploited this defense mechanism by using strain specific DMTase enzymes to pretreat DNA before electroporation or natural competence to improve transformation efficiency^58^. In this study, to identify potential *Fusobacterium* DMTases we could use to bypass R-M systems to increase the efficiency of transformation and DNA recombination, we queried the online databases REBASE^59^, FusoPortal^61^, and NCBI^62^ to characterize R-M systems. We analyzed 25 strains of *Fusobacterium nucleatum* in REBASE covering the subspecies *nucleatum (Fnn), animalis (Fna), vincentii (Fnv)*, and *polymorphum (Fnp)* for the number and classification of their R-M systems as shown in **Figure 1B**. There was an overall propensity for *Fn* strains to have a higher number of Type II DMTase genes, yet there was not a strong overall pattern of the number or class of R-M systems that differentiated the subspecies. As shown in **Figure 1C**, we highlight three strains of *Fn* covering subspecies *nucleatum* and *animalis*. The genetically tractable strain *Fnn* 23726 encodes 4 R-M systems as shown in **Figure 1C**; one Type I, two Type II, and one BREX system. *Fnn* 25586 lacks Type I R-M systems but has three Type II and two Type III DMTases that proved critical for enabling molecular genetics in this strain. Surprisingly, an extreme number of R-M systems were identified in *F. nucleatum* subsp. *animalis* 7_1 (*Fna* 7_1), for a total of 11 R-M systems (two Type I and nine Type II).

**Figure 1.**
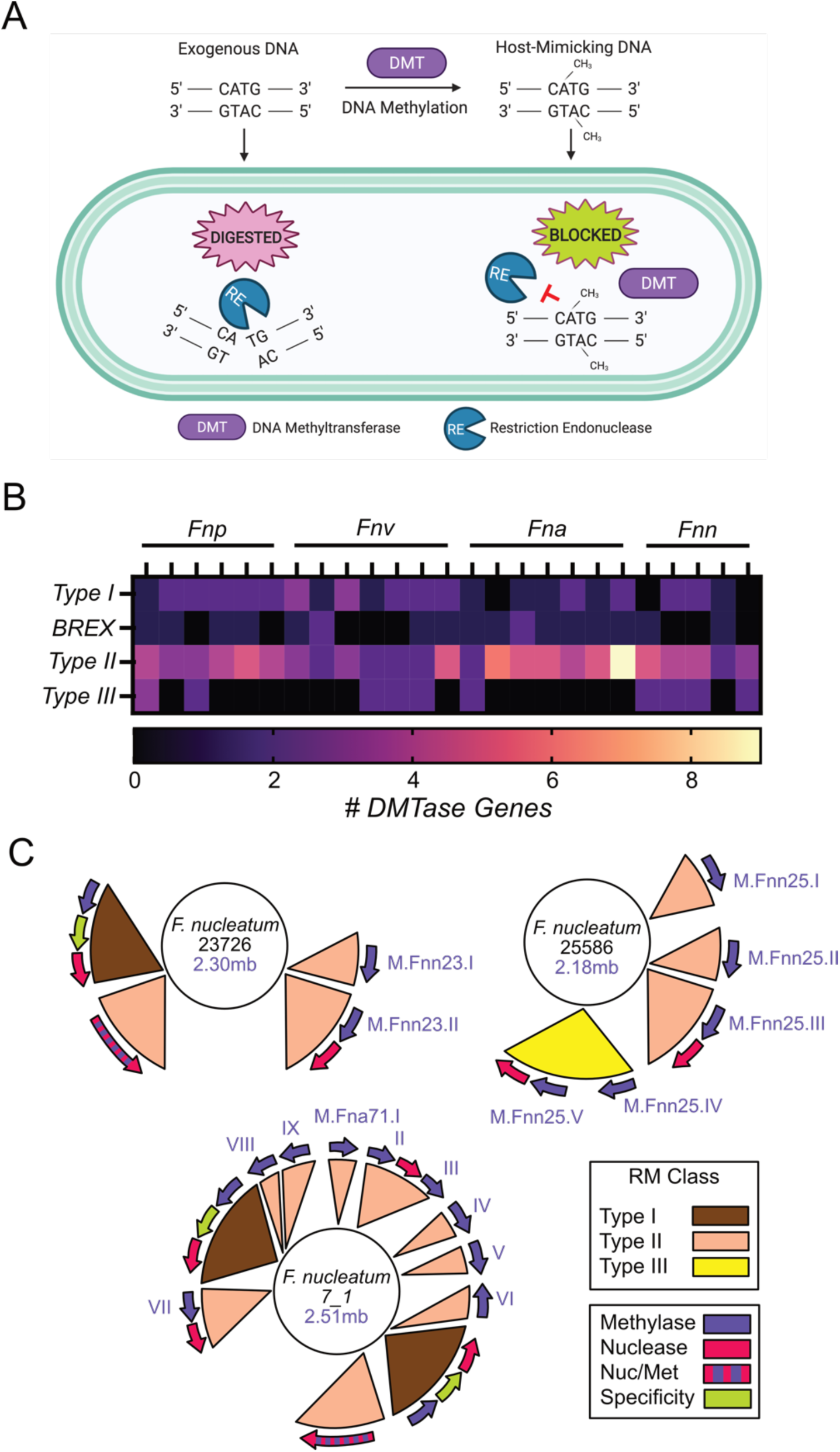
Restriction modification system classification in *Fusobacterium*. **(A)** Overview of how R-M systems utilize bacteria specific DNA methylation to mark the chromosome as ‘self’ DNA, thereby restriction digesting invading DNA that does not contain the proper methylation patterns. **(B)** Classification and quantitation of R-M systems in 25 strains of *Fn* covering the four subspecies: *polymorphum* (*Fnp*), *vincentii* (*Fnv*), *animalis* (*Fna*), and *nucleatum* (*Fnn*). **(C)** Genome location and renaming of Type II and Type III DMTases in three strains of *Fn* used in this study recreated from that on the REBASE website.

An orthodox Type II R-M system includes two independent genes in an operon: a DMTase and a REase. However, as shown in **Figure 1C**, the strong presence of lone methyltransferases was discovered in multiple *Fn* strains, and we later show these are crucial for protecting DNA for safe passage and genetics. These bioinformatic studies also confirmed the presence of the Type II BREX system in several *Fusobacterium* strains. The BREX system is generally composed of a 4-8 gene cluster,^12^ and in *Fusobacterium* is predicted to methylate adenine residues similar to *E*. coli^63^. However, since the restriction site for this enzyme is yet to be characterized, and these systems have not been shown to be important for efficient molecular microbiology efforts, we did not focus on using these enzymes for plasmid protection. Finally, no Type IV R-M systems were discovered in the *Fn* strains analyzed in this study. Utilizing REBASE, we identified the predicted DNA recognition and methylation sites for all Type II and Type III DMTases in the five strains of *Fn* that we use in this study: *Fnn* 23726, *Fnn* 25586, *Fna* 4_8, *Fna* 7_1, *Fnp* 10953 (**Table S1**). Nearly all DMTases are predicted to be adenine DNA methyltransferases, where methylation occurs at the nitrogen at position six in the ring (N^6^) of the adenine (N^6^-mA or 6mA), which is a common theme for A-T rich bacterial genomes (Fn >70% A-T).

### Recombinant production and characterization of DMTases

To focus our study, we chose to utilize and characterize all Type II and Type III DMTase enzymes in the strains *Fnn* 23726 and *Fnn* 25586. As shown in **Figure 2**, we cloned (**Fig 2A**), expressed, and purified (**Fig 2B**) five enzymes (M.Fnn23.I, M.Fnn23.II, M.Fnn25.I, M.Fnn25.IV, M.Fnn25.V). M.Fnn23.I and M.Fnn23.II were used to treat the plasmid pDJSVT13 as described below that we previously used to knock out the *galKT* genes in *Fnn* 23726^64^.

**Figure 2.**
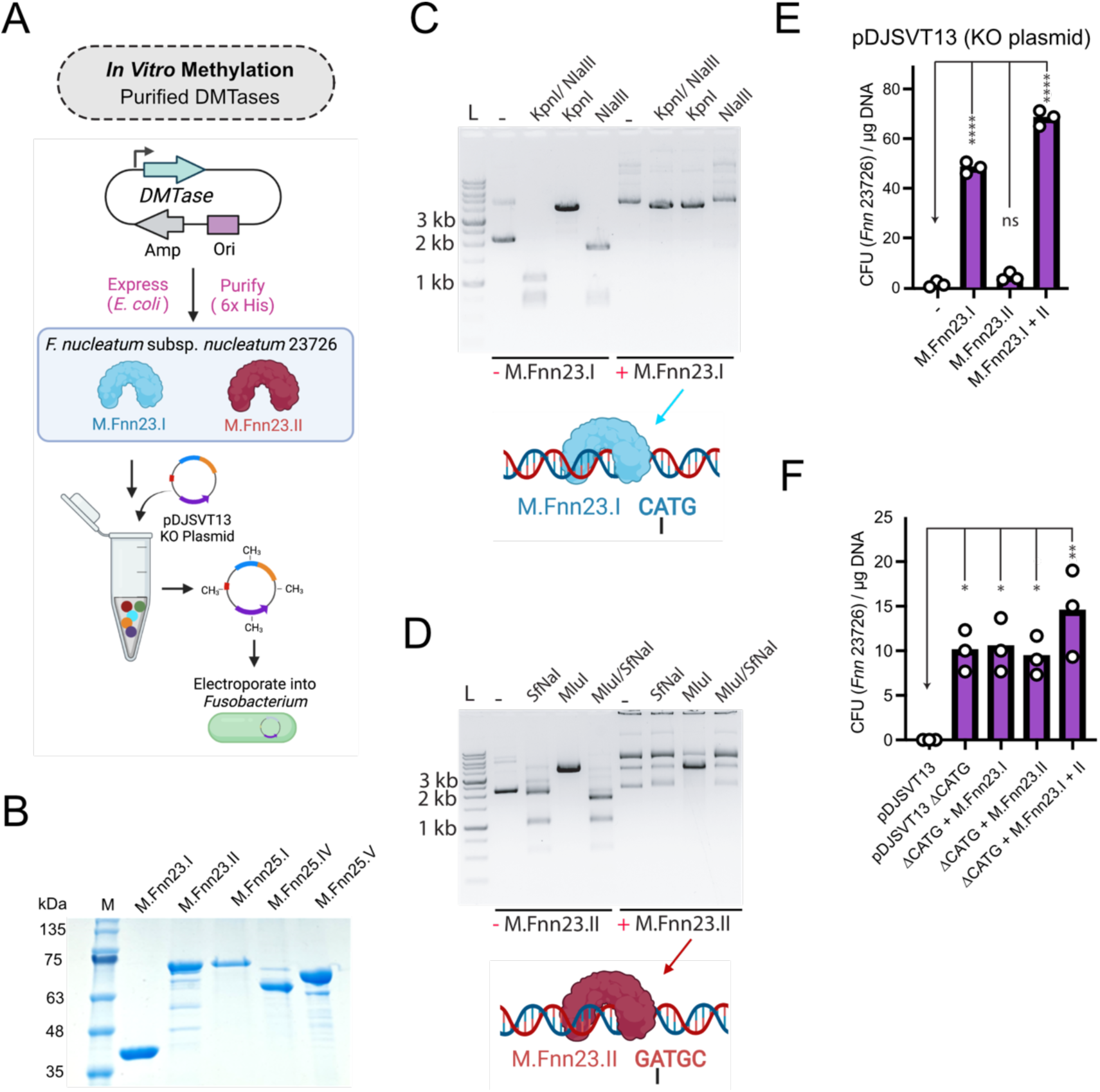
*Fn* DMTases protect plasmid DNA and allow for more efficient chromosomal plasmid incorporation in *Fnn* 23726. **(A)** Schematic of our process to produce recombinant DMTases that are next used to treat plasmid DNA *in vitro* prior to electroporation into *Fnn* 23726. **(B)** SDS-PAGE gel of purified of five purified DMTases from *Fnn* 23726 and *Fnn* 25586. **(C)** Methylation of pDJSVT13 with M.Fnn23.I protects against DNA cleavage by the REase NlaIII (NEB), which cuts at CATG sites. **(D)** Methylation of pDJSVT13 with M.Fnn23.II protects against DNA cleavage by the REase SfNaI (NEB), which cuts at GATGC sites. **(E)** Methylation of pDJSVT13 results in significantly more transformation and chromosomal incorporation. **(F)** By changing the CATG sequences to CACG, which are the target for the DMTase M.Fnn23.I, transformation efficiency is significantly increased even in the absence of methylation. Statistical values are as follows: ^ns^P >0.05, *P < 0.05, **P < 0.01, ***P < 0.001, ****P < 0.0001.

### Recombinant DMTases protect plasmid DNA from REase digestion

To show that our recombinant enzymes from *Fnn* 23726 were active, we identified commercially available REases that matched the methylation sequences of M.Fnn23.I and M.Fnn23.II. By methylating the plasmid pDJVST13 with M.Fnn23.I, we show that adenine methylation in the sequence CATG blocks cleavage by the endonuclease NlaIII, which recognizes the same sequence and cleaves 3’ to the guanine (**Fig 2C**). Next, we methylated pDJSVT13 with M.Fnn23.II, and show that methylation of the adenine in GATGC protects DNA from cleavage by SfNaI, which recognizes GCATC(N_5_) and cleaves 3’ to the N5 sequence (**Fig 2D**). This protection of DNA from cleavage by methylation indicates using these enzymes in tandem would allow more efficient homologous recombination in *Fnn* 23726 post electroporation.

### Plasmid DNA methylated with recombinant DMTases increases chromosomal integration for the *galKT* gene knockout plasmid pDJSVT13 in *Fn* 23726

As shown in **Figure 2E**, methylation of pDJSVT13 with M.Fnn23.I results in significantly more colonies after transformation, indicating protected DNA was not degraded before homologous recombination with the *galKT* operon in *Fnn* 23726. M.Fnn23.II alone did not have a drastic effect but did increase efficiency. Last, the combination of M.Fnn23.I and M.Fnn23.II resulted in the most robust increase in transformation and chromosomal incorporation, thereby greatly enhancing the efficiency of creating gene knockouts.

As M.Fnn23.I appears to be the dominant enzyme for protecting DNA in *Fnn* 23726, we made a pDJSVT13 ΔCATG plasmid, now called pDJSVT21, in which the four sites were eliminated with silent single nucleotide mutations. **Figure 2F** shows that pDJSVT21 transforms significantly better than pDJSVT13. The addition of M.Fnn23.I or M.Fnn23.II individually did not increase transformation efficiency over pDJSVT21. However, the addition of both enzymes did, which could mean that these enzymes are methylating at more than their bioinformatically predicted sites.

### *In vivo* methylation of plasmids increases transformation of gene knockout and transposon plasmids

We next developed plasmids that place the *m*.*fnn23*.*I* and *m*.*fnn23*.*II* DMTase genes downstream of a strong constitutive ‘Anderson’ promoter (iGEM part BBa_J23101) and before a short terminator (iGEM part BBa_00014). Plasmid pDJSVT24 contains *m*.*fnn23*.*I*, pDJSVT25 contains *m*.*fnn23*.*II*, and pDJSVT26 contains both *m*.*fnn23*.*I* and *m*.*fnn23*.*II* (**Fig 3A**). TOP10 *E. coli* containing one of the aforementioned plasmids expressing *Fnn* 23726 DMTases were transformed with the *galKT* gene knockout plasmid pDJSVT13, followed by plasmid purification from overnight growths. Upon transformation of this mixed pool of plasmids into *Fnn* 23726 and selection on thiamphenicol containing media to select for chromosomal incorporation of pDJSVT13, we show that this simple method of plasmid methylation is effective at significantly increasing transformation rate. M.Fnn23.I alone results in a marginal increase in efficiency, but methylation by both enzymes significantly increases transformation rates by more than fifty-fold (**Fig 3B**). As Top10 *E. coli* do possess Dam+ and Dcm+ methylation systems, we also used methylation free *E. coli* ER2796^65^ and show that plasmids purified from both strains transformed at the same rate into *Fnn* 23726 when pDJSVT26 was present and expressing M.Fnn23.I and M.Fnn23.II **(Fig3C)**.

**Figure 3.**
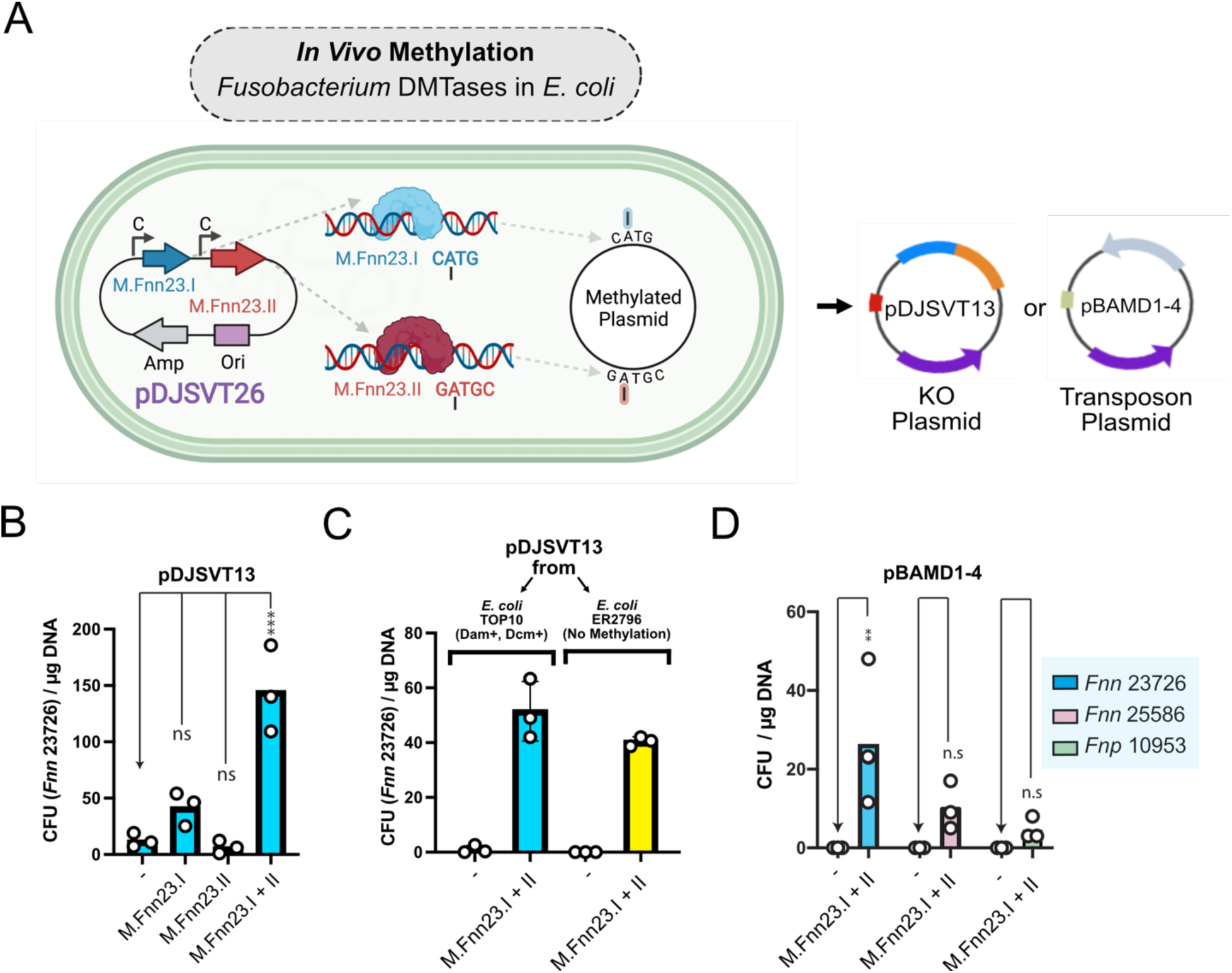
*In vivo* methylation in *E*.*coli* expressing M.Fnn23.I and M.Fnn23.II enhances plasmid transformation and chromosomal incorporation of plasmids and transposons. **(A)** Schematic of *in vivo* methylation of plasmids with *Fnn* 23726 DMTases. **(B)** Transformation of pDJSVT13 is significantly increase by co-expressing M.Fnn23.I and M.Fnn23.II. **(C)** Comparison of methylation positive (TOP10) and methylation negative (ER2796) *E. coli* reveals that native *E. coli* methylation does not inhibit the transformation of pDJSVT13 when *Fnn* 23726 DMTases are concurrently expressed. **(D)** *In vivo* methylation of the pBAMD1-4 transposon plasmid allows for transformation and chromosomal transposon insertion into multiple strains of *Fn*. Statistical values are as follows: ^ns^P >0.05, *P < 0.05, **P < 0.01, ***P < 0.001, ****P < 0.0001.

We next show that the mini Tn5 transposon harboring plasmid pBAMD1-4^66^ can be transformed into *Fnn* 23726, *Fnn* 25586, and *Fnp* 10953 after methylation with M.Fnn23.I and M.Fnn23.II, which we believe is the first time a spectinomycin resistant plasmid has been used for genetics in *Fusobacterium*. Important to note is that unmethylated plasmid was unsuccessful at producing transposon insertions in these three strains (**Fig 3D**). We do note that this system is not highly efficient and would benefit from using a more complete repertoire of DMTases from the respective strains. Overall, compared with *in vitro* plasmid treatment with recombinant DMTases, creation of an *E. coli* strain expressing *Fn* DMTases works just as well and requires less effort than purifying multiple proteins. However, the difficulty of creating plasmids with a significant number of DMTase genes makes this method increasingly challenging.

### Passaging of a plasmid in *Fn* allows for the transformation into additional strains

A common method of permitting plasmid to be transformed into a genetically recalcitrant strain of interest is to first transform into a similar, yet genetically competent strain, followed by repurification of the plasmid containing species specific methylation patterns (**Fig 4A**)^57^. This plasmid frequently can then be transformed into the strain of interest. Here we tested this classic method and show that passage of the episomal, multicopy *Fusobacterium* plasmid pHS30^49^ in *Fnn* 23726 can be purified and then transformed into *Fnp* 10953, but not *Fnn* 25586, *Fna* 7_1, or *Fna* 4_8. When plasmid is repurified from *Fnp* 10953, this plasmid can only be retransformed back into *Fnn* 23726, revealing that the RM systems in the other strains are not compatible with *Fnn* 23726 and *Fnp* 10953 (**Fig 4B)**. After methylating pHS30 with five DMTases to allow transformation into *Fnn* 25586, repurified plasmid was only able to be transformed into *Fnn* 23726. And once again, repurification of the plasmid from *Fnn* 23726 was only able to be transformed back into *Fnp* 10953.

**Figure 4.**
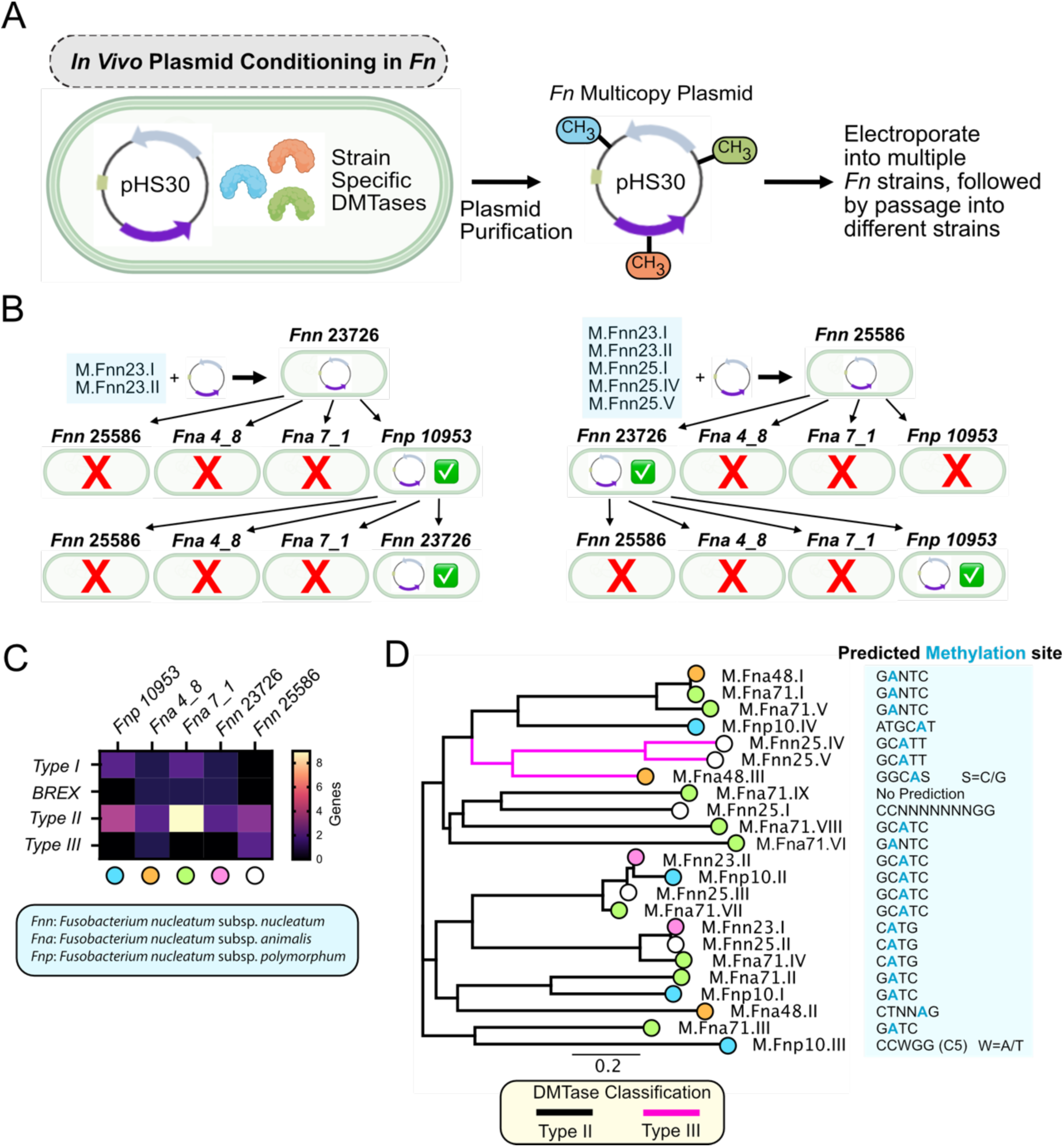
Passaging of a multicopy plasmid in *Fn* allows passage to additional strains. **(A)** Schematic of our passaging method for the *Fusobacterium* multi-copy plasmid pHS30, and purification of this plasmid for retransformation into different Fn strains. **(B)** pHS30 from *Fnn* 23726 can be transformed into *Fnp* 10953, and repurification from this strain allows transformation back into *Fnn* 23726. pHS30 from *Fnn* 25586 can be transformed into *Fnn* 23736, and repurification from this strain allows transformation back into *Fnp* 10953. **(C)** Heat map of the number of RM systems in the five *Fn* strains analyzed. Colored dots below the strain correlate with the strains of the enzymes found in the phylogenetic tree in Fig 4D. **(D)** Phylogenetic tree of 23 Class II and III DMTase genes from five *Fn* strains. Methylation sites as predicted by REBASE.

To better understand why there was limited plasmid passaging between *Fnn* strains, we analyzed the Type II and Type III DTMases in the five *Fn* strains tested above. We first compare the number of genes present in the strains for all classes of DMTases and note that all strains contain a higher number of Type II genes than the other classes (**Fig 4C**). However, other than strain *Fna* 7_1 having an extreme number of Type II RM systems, these data do not provide an obvious answer as to why the majority of these *Fusobacterium* strains are so genetically recalcitrant. To take a deeper look we assembled a phylogenetic tree of the 23 Type II and Type III DMTases from the five strains (**Fig 4D**). We have revealed clusters of enzymes with predicted DMTase recognition sites that could be exploited to produce a library of enzymes that could be used for bypassing RM systems in multiple strains. When analyzing the 23 DMTases from these five Fn strains, it stands out that the enzymes are predicted to methylate only ten recognition sites. These data also uncover that of the nine enzymes in *Fna* 7_1, which cover six predicted recognition sequences, only two of these sequences are predicted to be methylated by *Fnn* 23726 and *Fnn* 25586, leaving a large number of sequences unmethylated and the likely reason why plasmid was unable to be passed from these strains to *Fna* 7_1.

### *Fnn* 25586 and *Fnn* 23726 DMTases allow for the development of the first genetic system in *Fnn* 25586

*Fnn* 25586 is one of the classical strains that has been studied for more than four decades^67^, yet molecular studies have not been possible because of the inability to be transformed. Our goal was to use the same system we developed previously for gene knockouts in *Fnn* 23726^64^. As shown in **Figure 5** we used two DMTases from *Fnn* 23726 (M.Fnn23.I, M.Fnn23.II; same exact enzymes as M.Fnn25.II and M.Fnn25.III. **Fig 1C**), and three from *Fnn* 25586 (M.Fnn25.I, M.Fnn25.IV, M.Fnn25.V) to bypass RM systems in *Fnn* 25586 and create the first counterselectable genetic system. Purification of these recombinant enzymes was followed by methylation of pDJSVT13 and transformation by electroporation (**Fig 5A**). Colonies that grew on thiamphenicol containing plates indicated chromosomal integration by homologous recombination before (Fragment A) or after (Fragment B) the *galKT* operon (**Fig 5B**). PCR and sequencing verification of chromosomal integration (A or B single crossover; **Fig 5C-D**) was followed by double crossover events in non-selective media and plating on plates containing deoxygalactose, which verified excision of the *galKT* operon because the presence of *galKT* makes 2-Deoxy-D-galactose toxic (**Fig E-F**). *Fnn* 25586 *ΔgalKT* grows with the same fitness as Wild-Type *Fnn* 25586, WT *Fnn* 23726 and *Fnn* 23726 *ΔgalKT* (**Fig 5G**).

**Figure 5.**
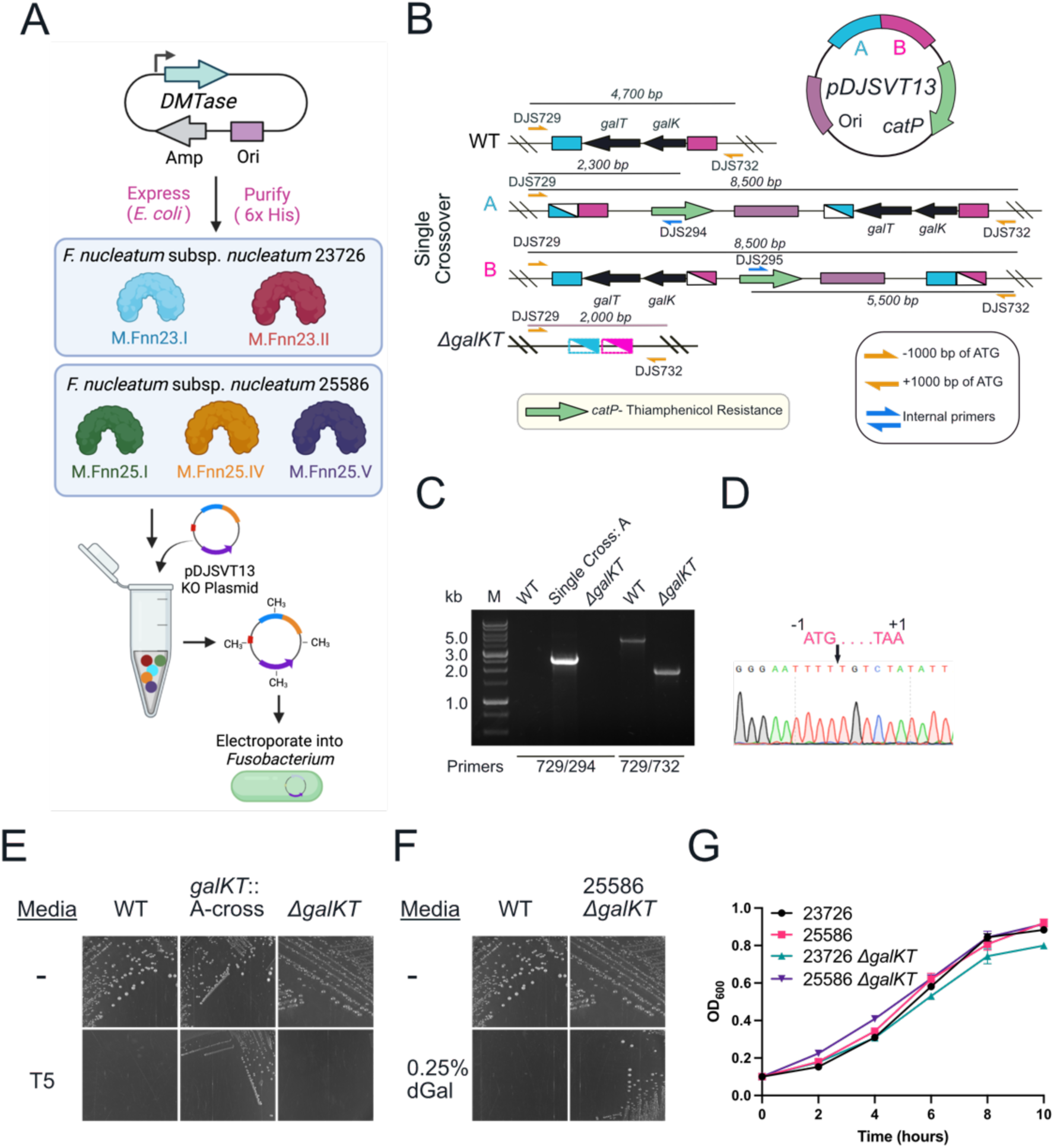
Development of a galactose selectable genetic system in *Fnn* 25586. **(A)** Schematic of the strategy to use five purified Fn DMTases to methylate plasmid pDJSVT13 to transform into *Fnn* 25586. **(B)** Schematic of single-crossover and *galKT* gene deletions using plasmid pDJSVT13, which first homologously recombines with up- and downstream sequences of the *galKT* operon. Primers noted that are used for PCR verification. **(C)** PCR verification of the initial chromosomal incorporation (A crossover) as well as the full operon deletion (*ΔgalKT*). **(D)** Sanger sequencing verification of a full, clean, deletion of the *galKT* operon. **(E)** Selection for A-crossover strains on thiamphenicol (T5) containing plates, and verification that the *ΔgalKT* strain has removed the vector and antibiotic cassette and no longer grows on thiamphenicol. **(F)** Proof of survival of *ΔgalKT* on plates containing deoxygalactose (dGal), which is toxic to wild type *Fnn* 25586. (G) Growth curves show no growth defect for *Fnn* 25586 *ΔgalKT* compared to WT *Fnn* 25586, WT *Fnn* 23726, and *Fnn* 23726 *ΔgalKT*.

### Development of *Fnn* 25586 *Δfap2* and *ΔfadA* strains

As a proof-of-concept, we made clean chromosomal deletions in genes *fap2* and *fadA* in *Fnn* 25586 *ΔgalKT* (**Fig 6**). This approach followed the same system that we initially used to knock out the *galKT* operon (**Fig 5**) to make a galactose selectable system possible. We report the clean deletions of the large, outer membrane, autotransporter adhesin *fap2* (> 10 kb) and the small, outer membrane adhesin *fadA* (390 bp), both of which have been studied extensively for their roles in *Fn* pathogenicity (**Fig 6A-F**)^68-71^. These gene deletions don’t cause any adverse growth phenotypes when compared to the parent strain *Fnn* 25586 *ΔgalKT* (**Fig 6G**). Our final experiment was to complement a gene deletion back onto the chromosome at the *arsB* gene^64^ **(Fig 6H)**, which confers arsenic resistance to bacteria but is not essential or necessary for *Fn* grown under laboratory conditions^72^. Because of this method, we witnessed that gene deletions in *Fnn* 25586 are now as efficient as *Fnn* 23726 (Data not shown), which has long been considered the most genetically tractable strain and therefore the strain with the most molecular studies. In addition, we report that like the system for *Fnn* 23726, there appear to be no differences in efficiency when deleting large (*fap2*; 10 kb) or small (*fadA*; 390 bp) genes in *Fnn* 25586.

**Figure 6.**
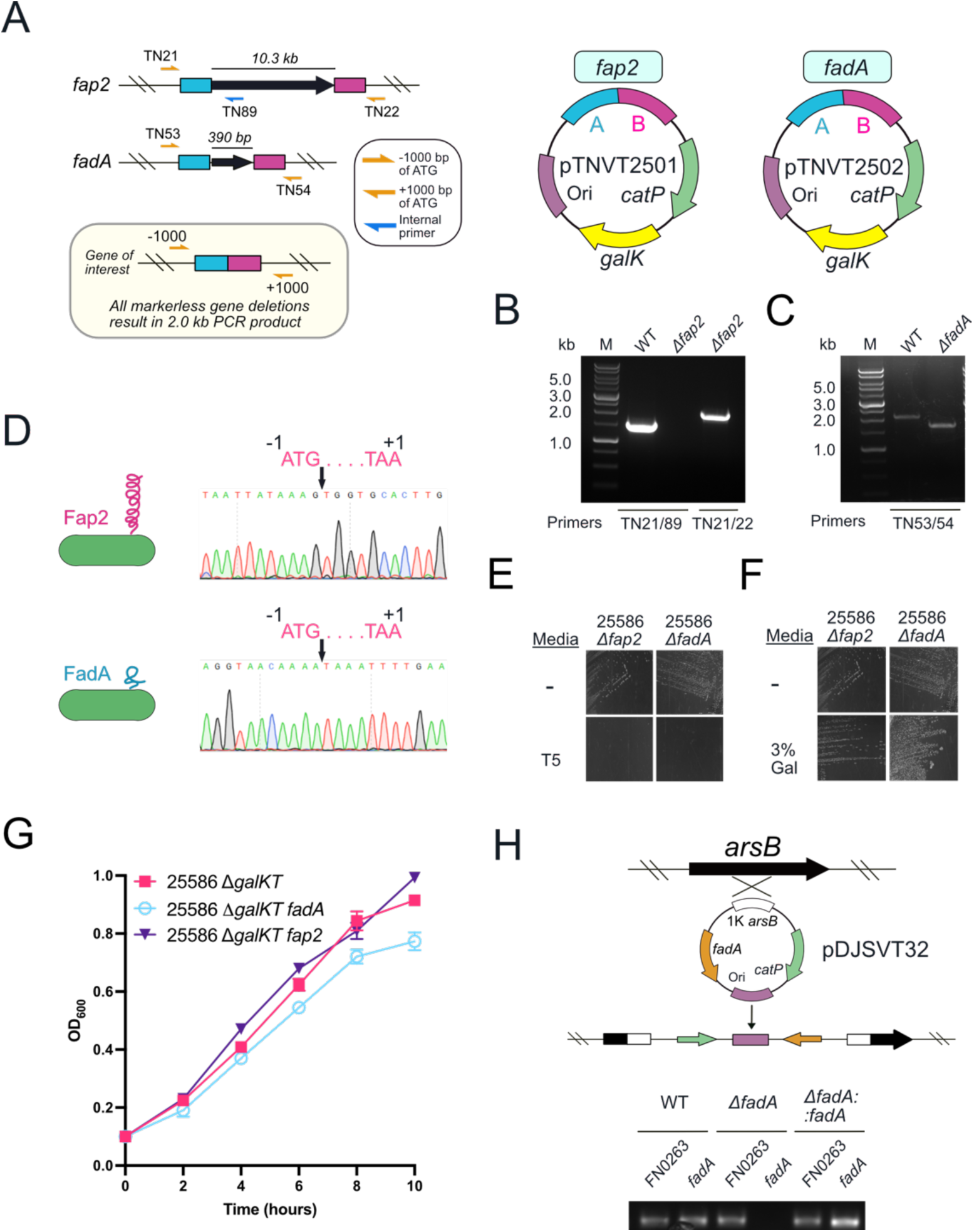
Gene deletions of *fap2* and *fadA*, as well as *fadA* complementation in *Fnn* 25586. **(A)** Schematic for the deletion of the genes *fap2* (>10kb) and *fadA* (390 bp) and primers used for PCR verification. Plasmids pTNVT01 and pTNVT02 correspond to plasmids created to delete *fap2* and *fadA*, respectively. **(B)** PCR verifying the *Δfap2* mutant in *Fnn* 25586. **(C)** PCR verifying the *ΔfadA* mutant in *Fnn* 25586. **(D)** Sanger sequencing verification of a full, clean, deletion of the *fap2* and *fadA* genes. **(E)** Streaking *Fnn* 25586 *Δfap2* and *ΔfadA* on thiamphenicol containing plates (T5) verifies the chromosomally integrated plasmid has been excised by homologous recombination. **(F)** Streaking *Fnn* 25586 *Δfap2* and *ΔfadA* on galactose containing plates (T5) verifies the chromosomally integrated plasmid has been excised by homologous recombination. **(G)** Growth curves show no fitness effects from *fap2* and *fadA* gene deletions. **(H)** Complementation of the *fadA* gene (*ΔfadA*::*fadA*) onto the chromosome at the *arsB* gene using a single-crossover homologous recombination plasmid and confirmation by PCR.

## DISCUSSION

Bacterial restriction-modification systems are important in both protection of bacteria from invading foreign DNA, as well as using methylation as an epigenetic switch to control gene regulation^73^. Our hypothesis was that if we could bypass *Fusobacterium* restriction modification systems it would enhance genetic efficiency in currently tractable strains, as well as leap the hurdle of developing new systems in strains that are inaccessible to molecular methods. Herein, we show that using strain specific DMTases from *Fusobacterium nucleatum* to methylate custom gene deletion plasmids leads to more efficient gene deletions, gene complementations on the chromosome, as well as the introduction of a multi-copy plasmid that could be used for a range of tasks including gene complementation and protein overexpression. Our results show a multifold increase in the efficiency of transformations and subsequent chromosomal incorporation of gene deletion plasmids in the genetically tractable strain *Fnn* 23726. To enhance genetics in this strain we cloned, expressed, and characterized two Type II DMTase enzymes which we renamed M.Fnn23.I and M.Fnn23.II. Using both *in vitro* and *in vivo* analysis, we verify that methylation of plasmid DNA blocks cleavage by the enzymes NlaIII and SfNaI, which cut at CATG and GATGC sites, respectively. We next show that each enzyme individually increases the efficiency of plasmid introduction but combining the two enzymes has a statistically significant affect.

We next set our focus on creating the first gene knockouts in *Fnn* 25586, which had not been accomplished in over forty years of studying the strain. To accomplish this, we produced five recombinantly expressed DMTases enzymes to treat plasmid DNA *in vitro*, followed by transformation by electroporation. Through this method we were able to create the first clean gene deletions and complementations in this strain, with deletions in *galKT, fap2, fadA*, and complementation of *fadA* back onto the chromosome. This markerless gene deletion system can produce an unlimited number of deletions in a single strain. Importantly, genetics in this strain are now as robust as that in *Fnn* 23726, which was previously the main strain used by most researchers due to its relative ease of use when compared to other non-transformable strains. We note that all five enzymes are necessary to protect plasmid DNA for safe passage into *Fnn* 25586; however, all DMTases that we produced and used in these studies retain their enzymatic activity after freeze-thaw cycles, making it a robust solution for researchers to implement.

We acknowledge that there are still major limitations to genetically modifying most strains of *Fusobacterium* because of their extreme differences in RM composition. Therefore, we understand that using a core combination of DMTase enzymes for universal protection across multiple *Fusobacterium* species may not be possible, as each strain frequently has unique DMTases that create a broad range of methylation patterns between strains. This has been reported before as Type II and Type III RM systems vary significantly even in evolutionarily similar strains of bacteria. We believe that future studies that combine DNA methylation analysis of DMTase deletion strains to identify exact methylation sequences with specific enzymes will lead to experimental determination of methylation sites by specific enzymes. To support this claim, previous studies have shown that using PacBio SMRTseq sequencing technology to determine the methylome of a bacterium results in the identification specific methylation sites, which can then be used to guide the creation of ‘Syngenic’ DNA plasmids that removes methylation and cleavage sites and therefore masquerades the DNA as self and is not cleaved by host^15^. An additional study used SMRTseq technology identified all DNA recognition sites and methylation patterns in multiple species of bacteria, followed by placing these sequences in a ‘methylation cassette’ within a plasmid that was then incubated with purified enzymes to identify specific methylation patterns^74^. Using this technique for highly recalcitrant strains of *Fusobacterium* would allow for the first true matching of methylation sites with *Fn* DMTases outside of bioinformatic predictions. Finally, one advantage we believe using recombinant DMTases has over this approach is that DNA methylation analysis and synthetic DNA based plasmids do not need to be made for each strain, which can keep down costs. However, one could argue that the effort of cloning and purifying the DMTases is on par with other methods of bypassing RM systems. Ultimately, we believe these methods are complementary and can be used in combination to enhance the chances of genetic modification in highly recalcitrant strains of *Fn*.

Potential future studies include an investigation into the role of using the Type I DMTase systems to methylate plasmids. In this area we briefly tried to recombinantly express HsdM and HsdS from *Fnn* 23726, but had difficulty achieving a pure, soluble protein complex. In addition, we report that we tried to use the Type I restriction modification inhibitor (Lucigen) in our transformations of *Fnn* 23726, but this did not change the transformation efficiency (data not shown). On a final note of the potential contribution of Type I RM systems, *Fnn* 25586 has no Type I systems and was still genetically impenetrable until using Type II and Type III enzymes. However, many transformable strains of bacteria have been made *hsdRMS* negative, which should be considered in the future as a method to potentially make more efficiently transformable strains of *Fusobacterium*.

Potential future strategies to increase genetic efficiency would be to delete the known REases in target strains. One disadvantage of this is the need to first transform and create a genetic system to be able to subsequently knock these genes out. But once accomplished an REase free strain would potentially bypass the need to treat entering plasmid DNA with DMTases. However, many of the Type II DMTases do not have a paired REases as shown in **Figure 1C**, therefore it is difficult to understand what could be cleaving the unmethylated DNA sequences that corresponds to specific enzymes. In addition, expanding beyond the realm of only studying *F. nucleatum* to other species including *F. necrophorum* could be key to understanding the pathogenicity of this species in Lemierre’s syndrome in humans^75^, as well as serious organ infections in livestock^76^.

In conclusion, we report that *Fn* DMTases can be used to methylate plasmid DNA, which then allows for efficient transformation and gene deletion in a well-studied strain, as well as a previously unmodifiable strain. The broader implications of this work are the enhanced ability to study the role of specific genes and corresponding virulence factors expressed by *Fn* during infection and disease. The methods in this study can be directly applied to target strains of interest within the scientific community, and therefore provides a roadmap for discovery biology that could lead to better understanding of how to inhibit the disease driving mechanisms of this oral, opportunistic pathogen.

## MATERIALS AND METHODS

### Bacterial strains and plasmids

All *E. coli* strains utilized in these studies were grown aerobically overnight at 37°C on solid Luria Bertani agar plates (10 g/L NaCl, 5 g/L tryptone, 10 g/L yeast extract) or in liquid Luria Bertani media. *Fusobacterium* strains were grown on solid agar plates made with Columbia Broth (Gibco), supplemented with hemin (5 μg/mL), menadione (0.5 μg/mL) and resazurin(1 μL/mL) under anaerobic conditions (90% N_2_, 5% H_2_, 5% CO_2_) at 37°C (Designated CBHK media). Liquid growths were inoculated from single *F. nucleatum* colonies and grown in CBHK liquid media under anaerobic conditions. Where necessary, antibiotics were supplemented at the suggested concentrations: gentamicin, 20 μg/mL; carbenicillin, 100 μg/mL; chloramphenicol, 10 or 25 μg/mL; thiamphenicol, 5 μg/mL (CBHK plates); and streptomycin 50 μg/ml (CBHK plates). The plasmids and bacterial strains utilized in these experiments are listed in **Table S2** and **Table S3**, respectively.

### Identification and classification of *Fn* DNA Methyltransferases

REBASE, a curated database of restriction enzymes, was used to identify the DNA methyltransferases present in the *F. nucleatum* subsp. *nucleatum* ATCC 23726 (GCA_003019875.1), *F. nucleatum* subsp. *nucleatum* ATCC 25586 (GCA_003019295.1), *F. nucleatum* subsp. *animalis* 7_1 (GCA_000158275.2), *F. nucleatum* subsp. *animalis* 4_8 (GCA_000400875.1), and *F. nucleatum* subsp. *polymorphum* 10953 (GCA_000153625.1) from the NCBI database. Type II and Type III DMTases were further bioinformatically characterized using NIH SMARTBLAST and pHMMER. SMARTBLAST and pHMMER provided conserved domains indicating function of DMTases. Phylogenetic analysis of *Fn* DMTase genes identified in REBASE were downloaded from NCBI and the NCBI identification numbers are supplied in **Table S1**. The tree and analysis were done in Geneious Prime 2022.1.1 using the Geneious Tree Builder function.

### Cloning, expression, and purification of DMTases

The DMTases M.Fnn23.I, M.Fnn23.II, M.Fnn25.I, M.Fnn25.IV, and M.Fnn25.V were cloned into pET16b under the control of an IPTG induced promoter for purification of the recombinant proteins using the C-terminal 6xHis tag and bench top metal affinity chromatography. In addition, M.Fnn23.I, M.Fnn23.II were cloned under the control of a constitutive promoter for continual expression in TOP10 *E. coli* to drive *in vivo* methylation of plasmids. All plasmids utilized and created in these studies are described in **Table S2** along with the bacterial strains in **Table S3** and primers in **Table S4**. The primers to clone the DNA methyltransferases were all ordered from Integrated DNA Technologies (IDT). For M.Fnn23.I and M.Fnn23.II, all constructs were made with *E. coli* codon optimized synthetic DNA was used for PCR. For DMTases from *Fnn* ATCC 25586, PCR was run with genomic DNA that was prepared with Wizard Genomic DNA Purification Kits (Promega).

Genes were amplified by PCR, and products were purified utilizing a PCR purification kit (Biobasic) and digested for 2 hours at 37 °C along with pET16b which was used as the expression vector and was obtained through EZ-10 Spin Column Plasmid Miniprep (Biobasic) with the restriction enzymes listed in **Table 4** with their respective primers. The vector was then dephosphorylated with Antarctic phosphatase (FastAP, Thermo Fisher Scientific) for 1 hour at 37 °C. Digested products were purified utilizing a spin column and ligated by T4 DNA ligase (New England Biolabs) for 1 hour at room temperature following manufacturer’s recommendations. Ligations were transformed into competent Mix&Go! (Zymo Research) Top10 *E. coli* and plated on LB solid agar plates supplemented with 100 μg/mL carbenicillin (ampicillin). Confirmation of positive clones was performed by digestion and if applicable positive clones were then transformed into ARTIC(DE3) RIL or LOBSTR-BL21(DE3) RIL^77^ for recombinant protein expression.

For protein expression *E. coli* cells were grown in LB (15g/L NaCl, 15 g/L tryptone, 10g/L yeast extract) medium at 37°C, 250 rpm shaking until OD=0.6. At OD=0.6. cells were induced with 50 μM Isopropyl β-d-1-thiogalactopyranoside (IPTG) (GoldBio). Expression was carried out at 8 °C and cells were collected at 20 hours after inoculation by centrifugation at 5000×g for 20 min at 4 °C. Bacterial pellets were resuspended in a lysis buffer (20 mM Tris, pH 7.5, 400 mM NaCl, 20 mM imidazole). Bacteria were lysed by an EmulsiFlex-C3 homogenizer (Avestin) at 10,000 kPa. Unlysed cells and insoluble material was separated by centrifugation at 15,000×g for 20 minutes at 4°C and then discarded. The supernatant containing the 6xHis-tagged DMTases was stirred with 6 mL of NiCl_2_-charged chelating Sepharose beads (GE Healthcare) for 30 minutes at 4°C. The column was washed with 400 mL of wash buffer (20 mM Tris, pH 7.5, 400 mM NaCl, 40 mM imidazole). After washing, the methyltransferases were eluted in 10 mL of elution buffer (20 mM Tris, pH 7.5, 400 mM NaCl, 250 mM imidazole). The purified protein was then directly put into dialysis in a buffer (20 mM Tris, pH 7.5, 150 mM NaCl, 10% glycerol). Protein concentrations were calculated using a Qubit fluorometer and BCA assays, followed by freezing at -80 °C for long-term storage.

### *In vitro* treatment of plasmid DNA with *Fusobacterium* DNA Methyltransferases

Plasmid DNA (35-40 μg), prepared from E.coli TOP10 using the EZ-10 Spin column plasmid DNA mini-prep from Biobasic, was combined in a 30 μL reaction with 160 μM SAM (New England Biolabs), 1X Cutsmart buffer (New England Biolabs) and 1 μM of one or more DMTases. The reaction mixes were incubated at 37°C for 2 hours and then plasmid was extracted by adding 1 volume of Phenol:Chloroform:Isoamyl Alcohol, 25:24:1 Mixture (bioWORLD) and vortexed for 20 seconds. Mixtures were then centrifuged at 16,000×g for 5 minutes. Plasmid DNA was precipitated and washed with ethanol and dissolved in ultrapure water (bioWORLD), followed by further purification Plasmid DNA was purified from overnight expression or co-expression was isolated with an alkaline lysis/column purification technique using the EZ-10 Spin Column Plasmid Miniprep (Biobasic). Plasmid DNA was further purified for use in electroporation by precipitation overnight at -80°C in 75% ethanol with sodium acetate (pH 5.5) and 0.1 μg/ml glycogen. After 3 hours minimum of incubation at -80°C sample was spun at 4°C for 30 minutes at 16,000×g to pellet the DNA and washed five times with 70% ethanol carefully by spinning at 14,000×g for 3 mins. Pellet was then dried at room temperature for 10–13 minutes. Finally, 15 μL of ultrapure water was added and incubated at 37°C for 1 hour to solubilize the pellet. DNA concentrations were determined using a NanoDrop spectrophotometer.

### Co-expression of plasmid DNA with *Fn* DMTases for *in vivo* methylation

Using the expression vector (constitutive activity) pET16b with the DNA methyltransferase under an Anderson medium promoter as described in **Table S2**, we methylated pDJSVT13 *in-vivo*. Both pET16b (Gene 622 and Gene 635) and pDJSVT13 were transformed into *E*.*coli* top10 and grown in LB (15 g/L NaCl, 15 g/L tryptone, 10g/L yeast extract) medium at 37°C, 250 rpm shaking for 24 hours.

### REase protection assays

Plasmid DNA (1 μg) prepared from *E*.*coli* TOP10 strain using the Biobasic mini-preparation procedure, was combined with Cutsmart buffer (New England Biolabs, 50 mM Potassium Acetate, 20 mM Tris-acetate, 10 mM Magnesium Acetate, 100 μg/mL BSA (pH 7.9)), 160 μM SAM (New England Biolabs), and 1 μM of the correspondent DMTases. As a control plasmid DNA (1 μg) was mock treated in reaction buffer without the methyltransferases. All samples were incubated 1 hour at 37 °C with the restriction enzymes, single cutters KpnI and MluI or predicted restriction sites NlaIII and SfaNI (New England Biolabs). For single-cut linearization, plasmid DNA was digested with restriction enzyme KpnI following manufacturer’s instructions (NEB). After two hours at 37°C the ultrapure DNA underwent phenol chloroform extraction and ethanol precipitation at -80°C as described previously for ultrapure DNA purification. Samples were analyzed in a 1% agarose gel with ethidium bromide and imaged on a Syngene G:Box imager as shown in **Figure 2C-D**.

### *Fn* transformation by electroporation

All *Fn* strains were competently prepared by inoculating and growing a 100-mL anaerobic culture in CBHK media to lag phase (A_600_ = 0.1) followed by centrifugation of bacteria at 3200×g for 10 minutes. The supernatant was removed, and the resulting pellet was washed three successive times utilizing 1 mL of ice-cold 20% glycerol in deionized H_2_O and 1mM MOPS at 14,000×g for 3 minutes. Bacterial pellet was then resuspended in a final volume of 80 μL of ice-cold 20% glycerol and 1mM MOPS. Bacteria were transferred to cold 1 mm (Lonza) electroporation cuvettes, and 3 μg (concentration >300 ng/μL) of plasmid was added before electroporating at 2.5 kV/cm, 50 μF, 360Ω, using a BTX Electro Cell Manipulator 600 (Harvard Apparatus). The electroporated cells were promptly transferred by syringe into a sterile, anaerobic tube with 4 mL of recovery medium (CBHK, 1 mM MgCl_2_) and incubated at 37 °C for 20 h with no shaking in an anaerobic chamber. After the recovery outgrowth, cells were centrifuged at 14,000×g for 3 minutes, supernatant was removed, and pellet cells were resuspended in 0.2 mL of recovery medium. Resuspension was plated on CBHK plates with 5 μg/mL thiamphenicol and incubated in an anaerobic 37 °C incubator for two days for colony growth. The transformation efficiency represents the number of thiamphenicol or streptomycin resistant colonies per microgram of DNA. Electroporation was conducted in triplicate as independent experiments.

### Utilizing plasmid methylation to enable a galactose-selectable gene deletion system in *Fnn* 25586

A galactose selectable gene deletion system for *Fnn* 23726 was previously developed in our lab and reported in detail in Casasanta et al^64^. As *Fnn* 23726 and *Fnn* 25586 are extremely similar at the DNA level, the plasmid pDJSVT13 that was previously used to delete the *galKT* operon in *Fnn* 23726 was also used on *Fnn* 25586 because of 100% nucleotide identity in the up and downstream regions cloned for homologous recombination and gene deletion. pDJSVT13 was conditioned with methylated with five DMTase enzymes (M.Fnn23.I, M.Fnn23.II, M.Fnn25.I, M.Fnn25.IV, M.Fnn25.V) using the same conditions as describe above for the in vitro methylation protocol. Ultrapure DNA (3 µg) was electroporated (2.5 kV, 50-µF capacitance, 360-Ω resistance, 0.2-cm cuvette) into competent *Fnn* 25586, and single chromosomal crossovers of the pDJSVT13 plasmid were selected for on thiamphenicol. Colonies were then inoculated into antibiotic free CBHK media overnight at 37 °C to allow for a second crossover event, which effectively deletes the target gene and also the remaining plasmid that was integrated into the chromosome. Next, 100 µL from this culture was streaked on solid medium containing 0.25% 2-deoxy-D-galactose to select for *galKT* gene deletions, as the absence of the *galT* gene makes 2-deoxy-D-galactose nontoxic to *Fnn. galKT* gene deletions were verified by PCR and sanger sequencing. This new strain, *Fnn* 25586 *ΔgalKT*, which we now name TNVT2501, is now the base strain used to create all future targeted gene deletions. Bacterial transformation of TNVT2501 allows for initial chromosomal integration and selection with thiamphenicol, followed by selection for double crossover gene deletions on solid medium containing 3% galactose. We have shown that deletion of the *galKT* operon in *Fnn* 25586 does not result in altered fitness.

### Creating *Fnn* 25586 *Δfap2* and *Fnn* 25586 *ΔfadA*

As a proof of concept, we next generated targeted gene deletions in the *Fnn* 25586 *ΔgalKT* background and in the two most well-studied *Fn* virulence factors: *fap2* and *fadA*. The first step is to use the plasmid pDJSVT7, which contains a FLAG::*galK* gene under the control of a *Fusobacterium necrophorum* promoter. Briefly, 750 bp directly upstream and downstream of the *fap2* and *fadA* genes were amplified by PCR and fused by OLE-PCR. PCR product was digested with KpnI/MluI ligated into pDJSVT7 digested with the same enzymes, followed by transformation into TOP10 *E. coli* and selection on LB plates containing chloramphenicol. Positive clones were identified by restriction digest and sanger sequencing to verify the new gene deletion plasmids pTNVT2501 (*fap2)* and pTNVT2502 (*fadA)* (**Fig 6A**). pTNVT2501 and pTNVT2502 were next electroporated (3 µg of DNA, 2.5 kV, 50-µF capacitance, 360-Ω resistance, 0.2-cm cuvette) into competent *Fnn* 25586 *ΔgalKT* and chromosomal integration was selected for on thiamphenicol (single chromosomal crossover), followed by selection on solid medium containing 3% galactose, which produces either complete gene deletions or wild-type bacteria revertants. Gene deletions were verified by PCR and Sanger sequencing as shown in **Figure 6**. The new strain names are TNVT02 and TNVT03 for the *Δfap2* and *ΔfadA* in *Fnn* 25586. We showed that this system was accurate down to the single base level for creating clean genome excisions that therefore allow for the deletion of an unlimited number of genes.

### Complementation of a *fadA* gene deletion in *Fnn* 25586

We previously created the gene complementation vector pDJSVT11 to create single-copy chromosomal complementation at a chromosomal location within the *arsB* gene^64^. Our previously developed plasmid pDJSVT32 was used to complement *Fnn* 23726 *ΔgalKT fadA* and was also used to complement *Fnn* 25586 *ΔgalKT fadA* (TNVT03). Briefly this plasmid contains a 1000 bp central region of the *arsB* gene, driving homologous recombination, which results in chromosomal insertion of the thiamphenicol resistance plasmid.

Complementation was selected for on CBHK plates containing thiamphenicol, followed by inoculation into liquid CHBK containing thiamphenicol. Complementation was further verified by PCR of the *fadA* gene as shown in **Figure 6H**.

### Statistical analysis

All statistical analysis was performed in GraphPad Prism Version 8.2.1. For single analysis, an unpaired Student’s t test was used. For grouped analyses, Two-way ANOVA was used. In each case, the following P values correspond to star symbols in figures: ^ns^P >0.05, *P < 0.05, **P < 0.01, ***P < 0.001, ****P < 0.0001. To obtain statistics, all studies were performed as three independent biological experiments. For all experiments in which statistical analysis was applied, an N of 3 independent experiments was used (details in figure legends).

## ACKNOWLEDGMENTS

This research was supported by the National Institutes of Health through an NCI R21 Award (grant no. 1R21CA238630-01A1 to D.J.S.), the College of Agriculture and Life Sciences at Virginia Tech (to D.J.S.), the Institute for Critical Technology and Applied Science at Virginia Tech (to D.J.S.) and the USDA National Institute of Food and Agriculture (to D.J.S.). Select figures were made with a paid subscription of Biorender.com

## CONFLICT OF INTEREST

The authors declare no conflict of interest

## AUTHOR CONTRIBUTIONS

A.U., T.T.D.N., and B.E.S. curated and analyzed the data, designed/optimized the methodology, wrote, reviewed, and edited the manuscript. K.J.W and B.W. curated the data and reviewed and edited the manuscript. D.J.S helped conceptualize, supervise, and acquire funding for the study, performed data analysis, curated, and analyzed the data, designed/optimized the methodology, wrote, reviewed, and edited the manuscript.

## DATA AVAILABILITY STATEMENT

Materials are available upon reasonable request with a material transfer agreement with Virginia Tech for bacterial strains, or through the Addgene repository for plasmids.

## SUPPLEMENTAL MATERIAL

**Figure S1.**
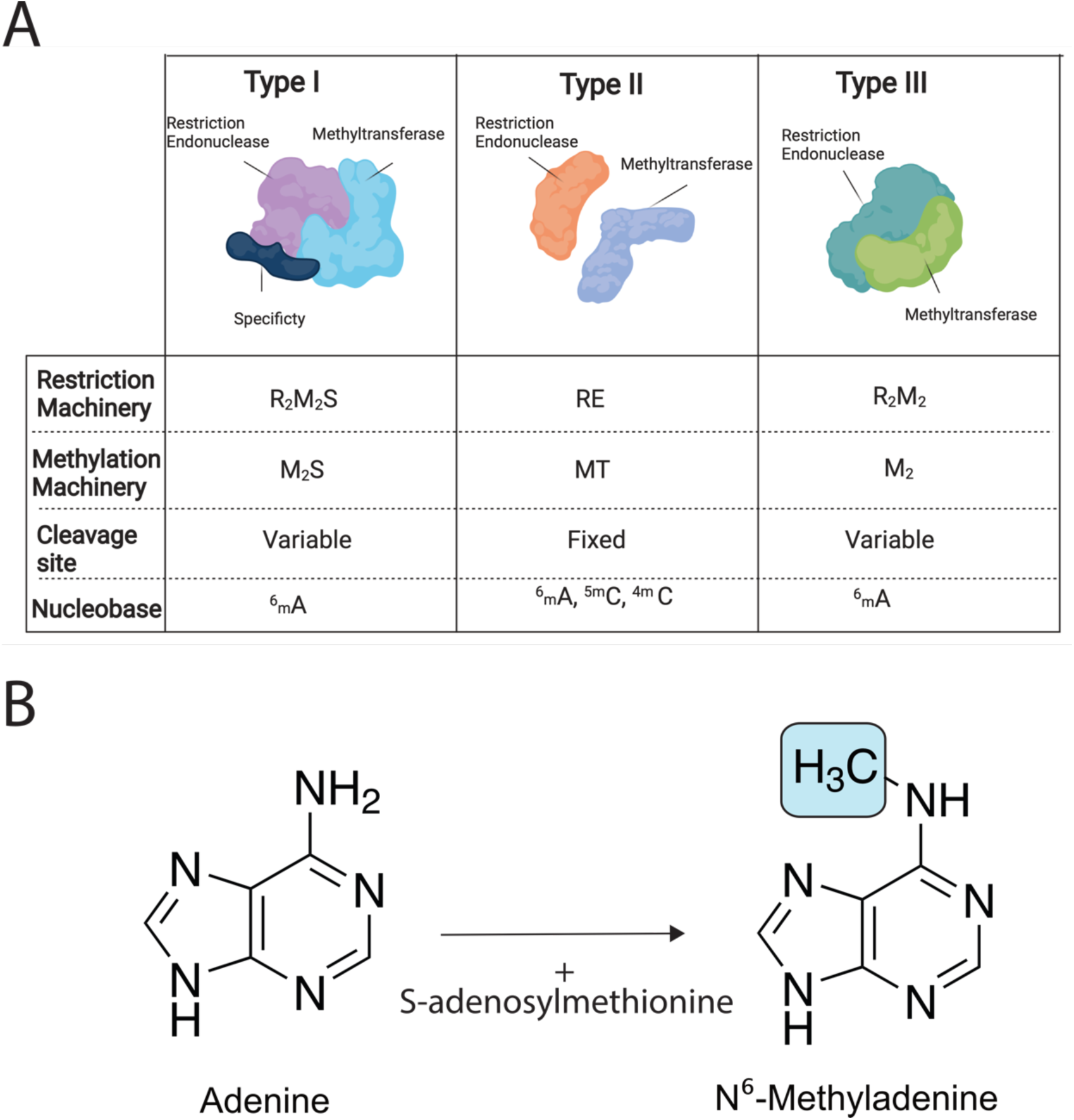
R-M systems and DNA methylation. **(A)** R-M systems are classified as Type I, Type II, and Type III according to their molecular structure, subunit composition, cleavage position, restriction site, and cofactor specification. **(B)** Nearly all methylation in *Fusobacterium* is predicted to be on adenine or adenosine residues within DNA and is added to nitrogen at the 6^th^ position to create N6-Methyladenine (N^6^-mA or ^6^mA).

**Table S1.**
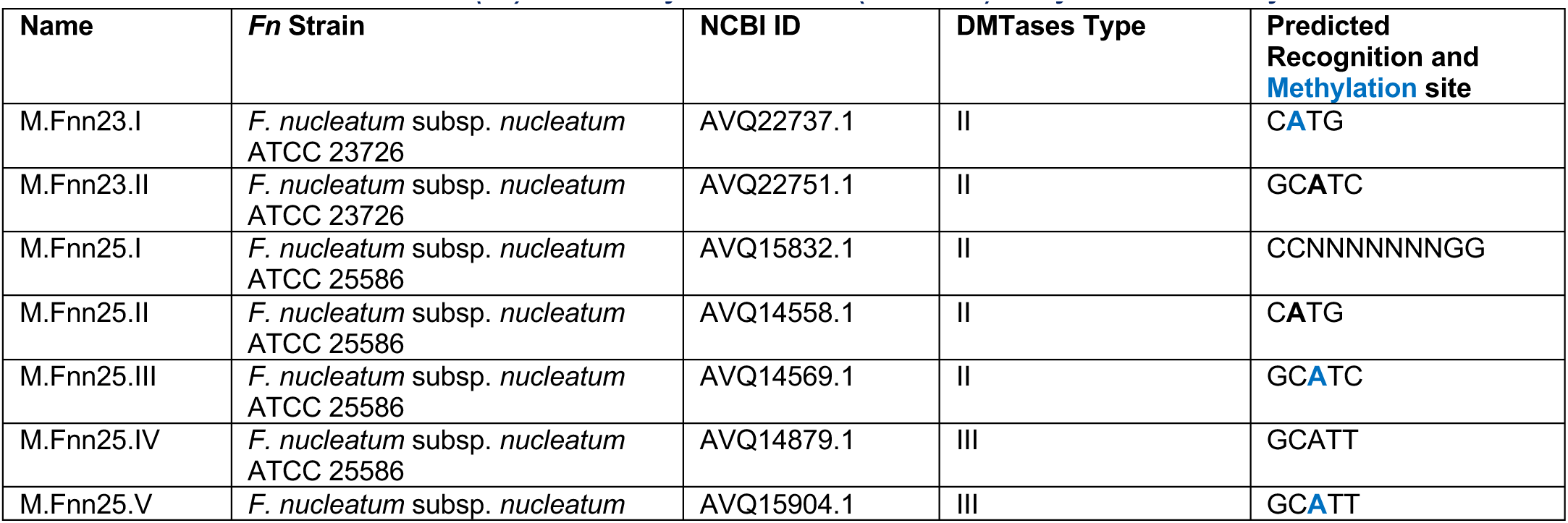
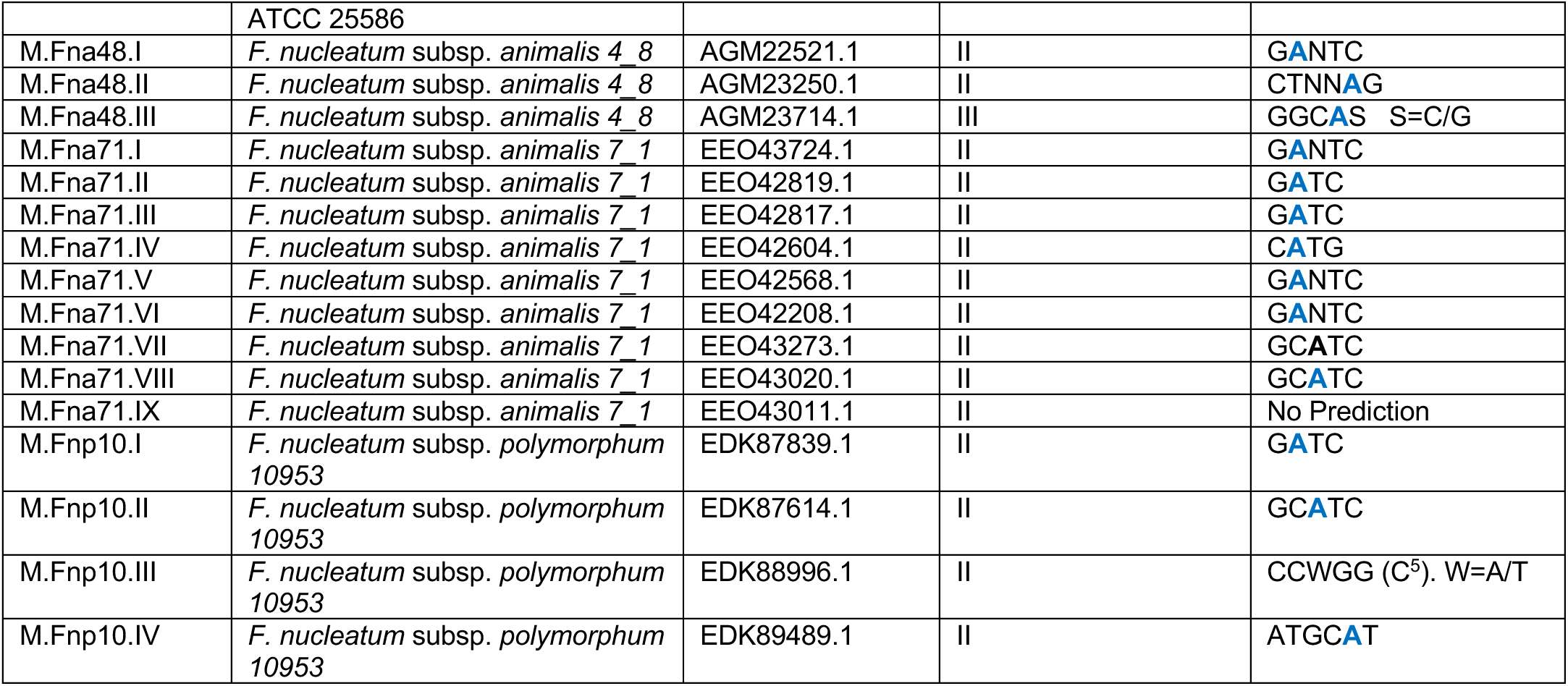
*Fusobacterium nucleatum (Fn)* DNA methyltransferases (DMTases) analyzed in this study.

**Table S2.**
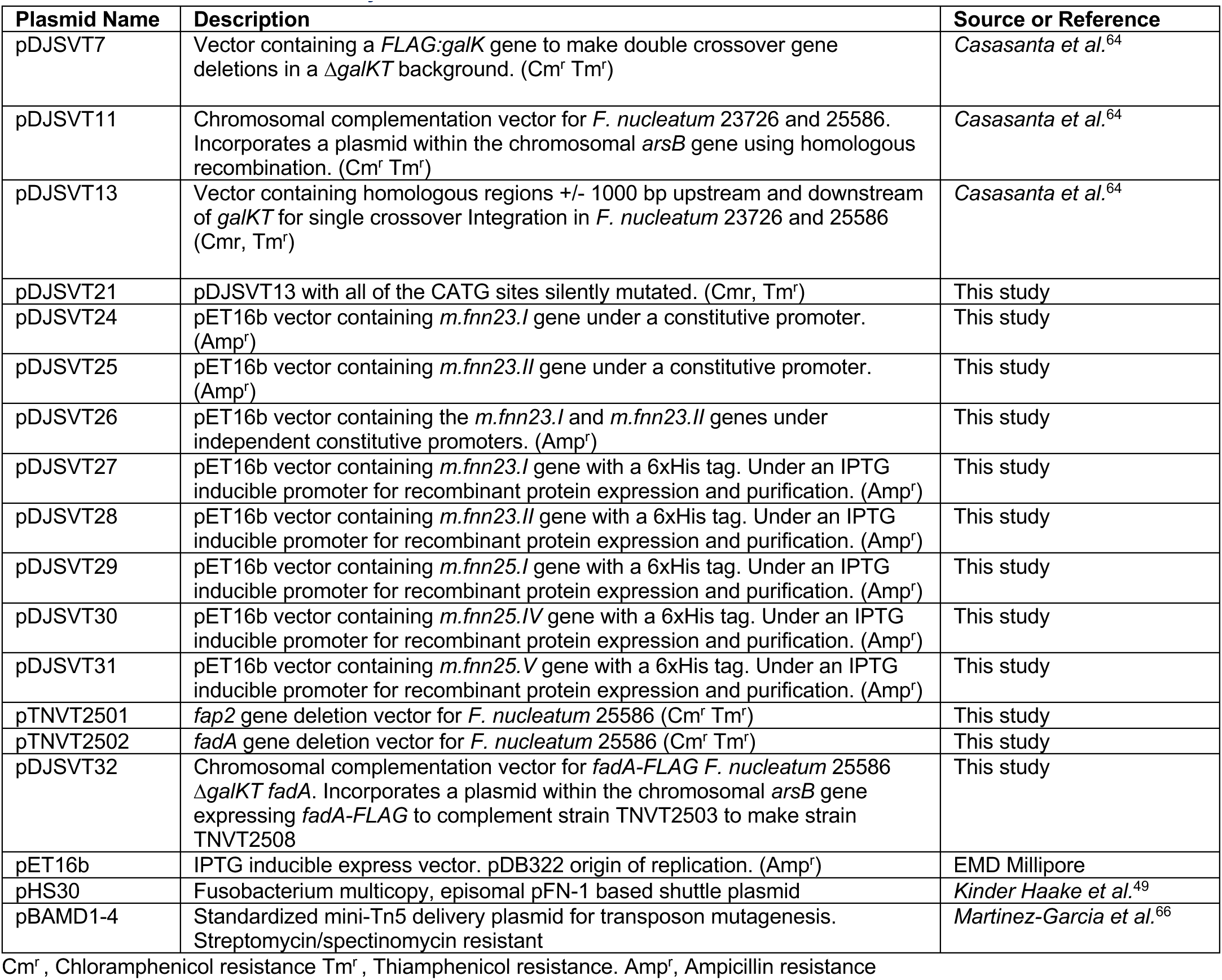
Plasmids used in this study.

**Table S3.**
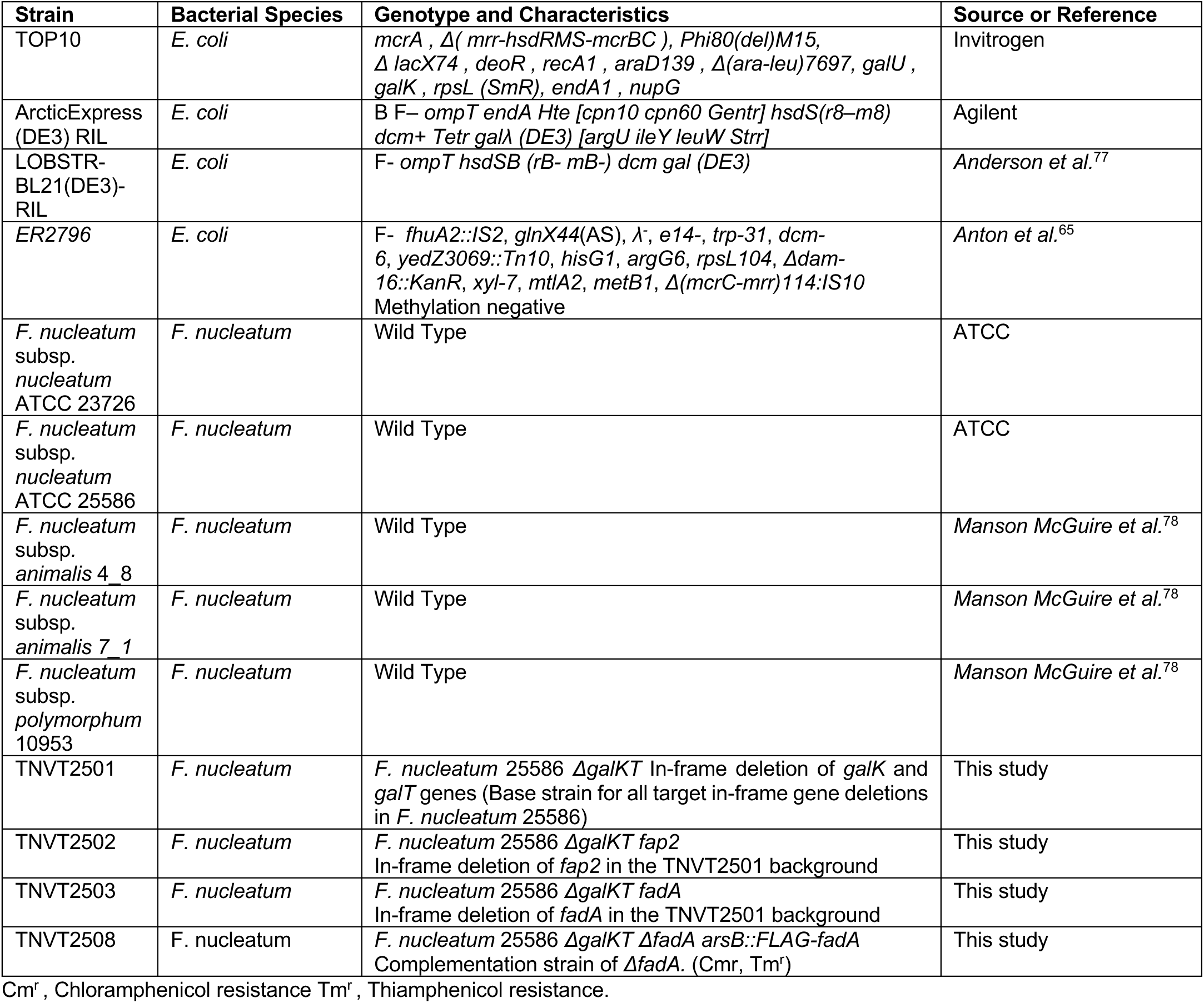
Bacterial strains used in this study.

**Table S4.**
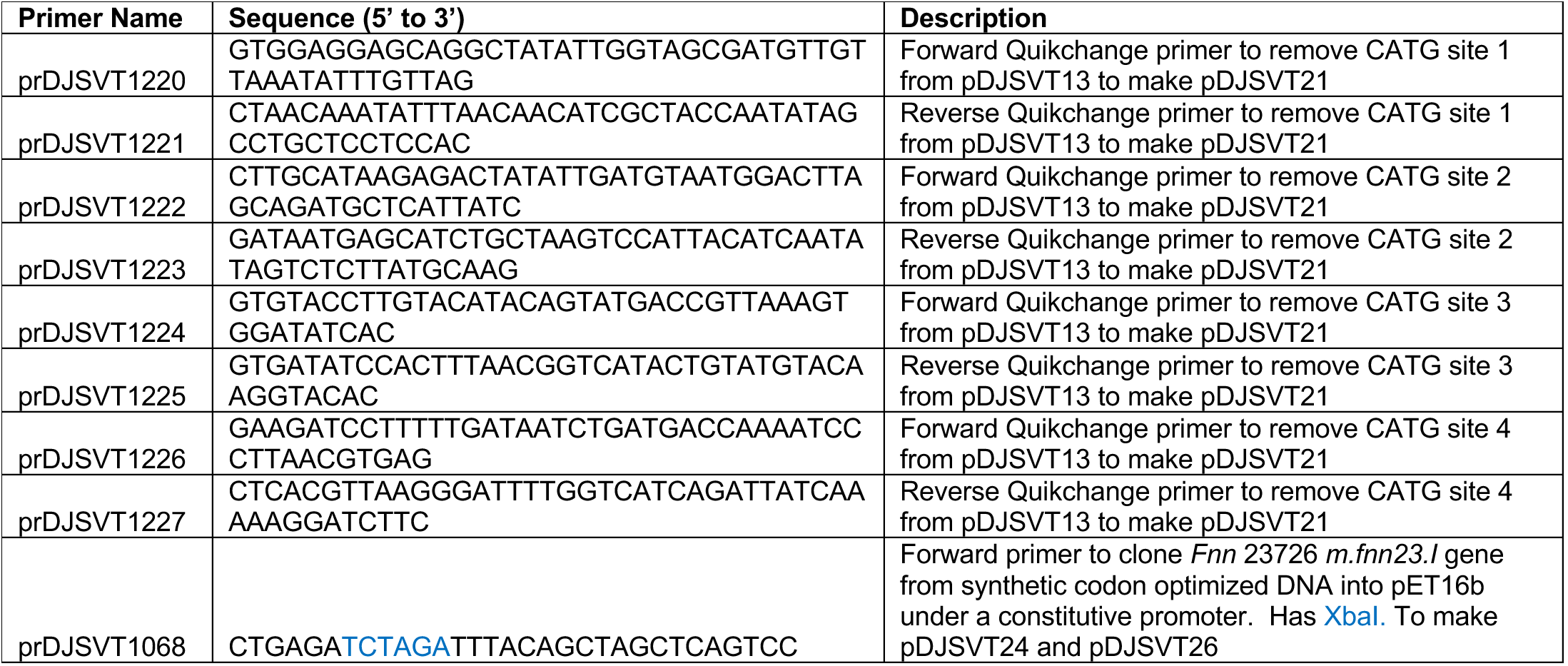
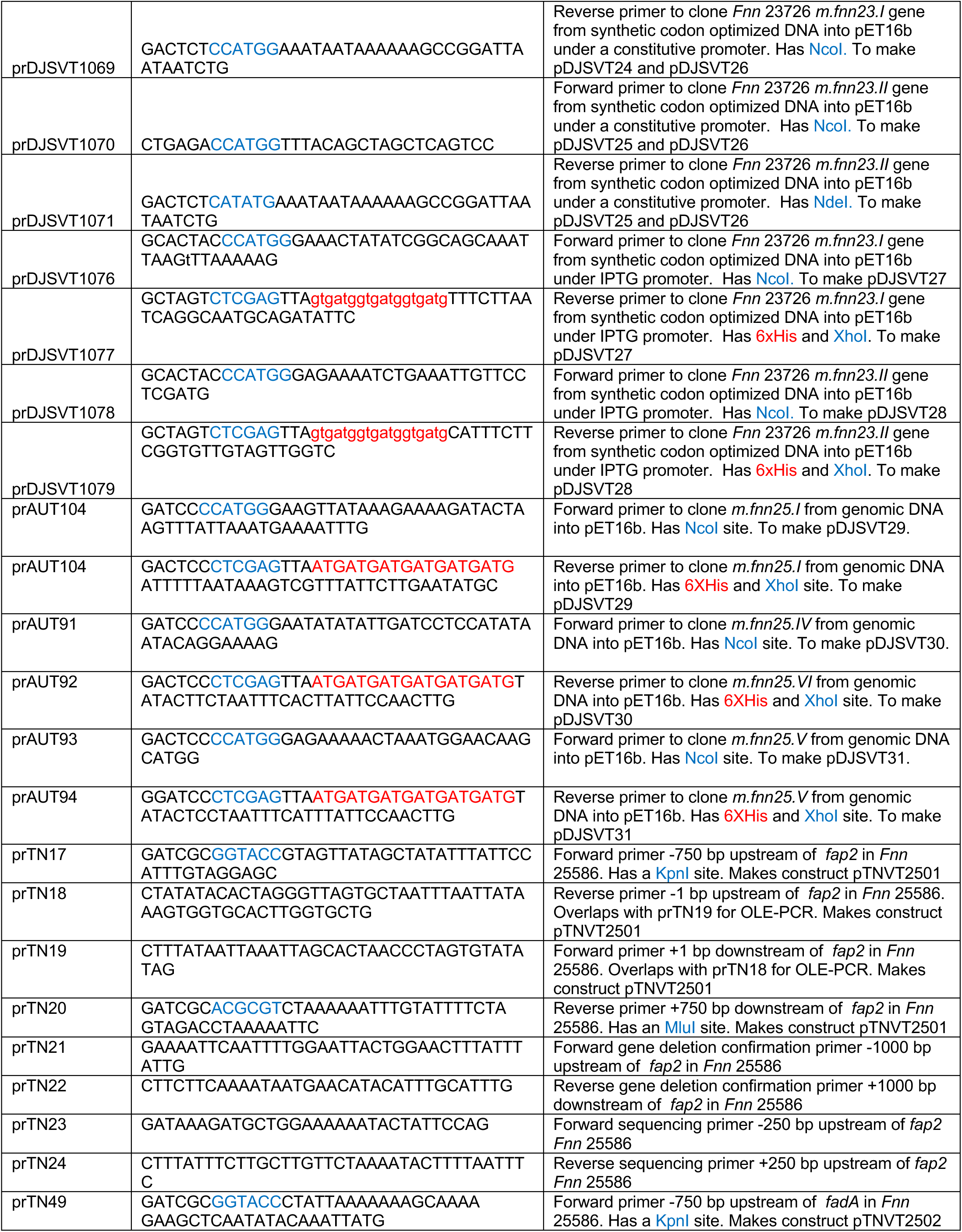
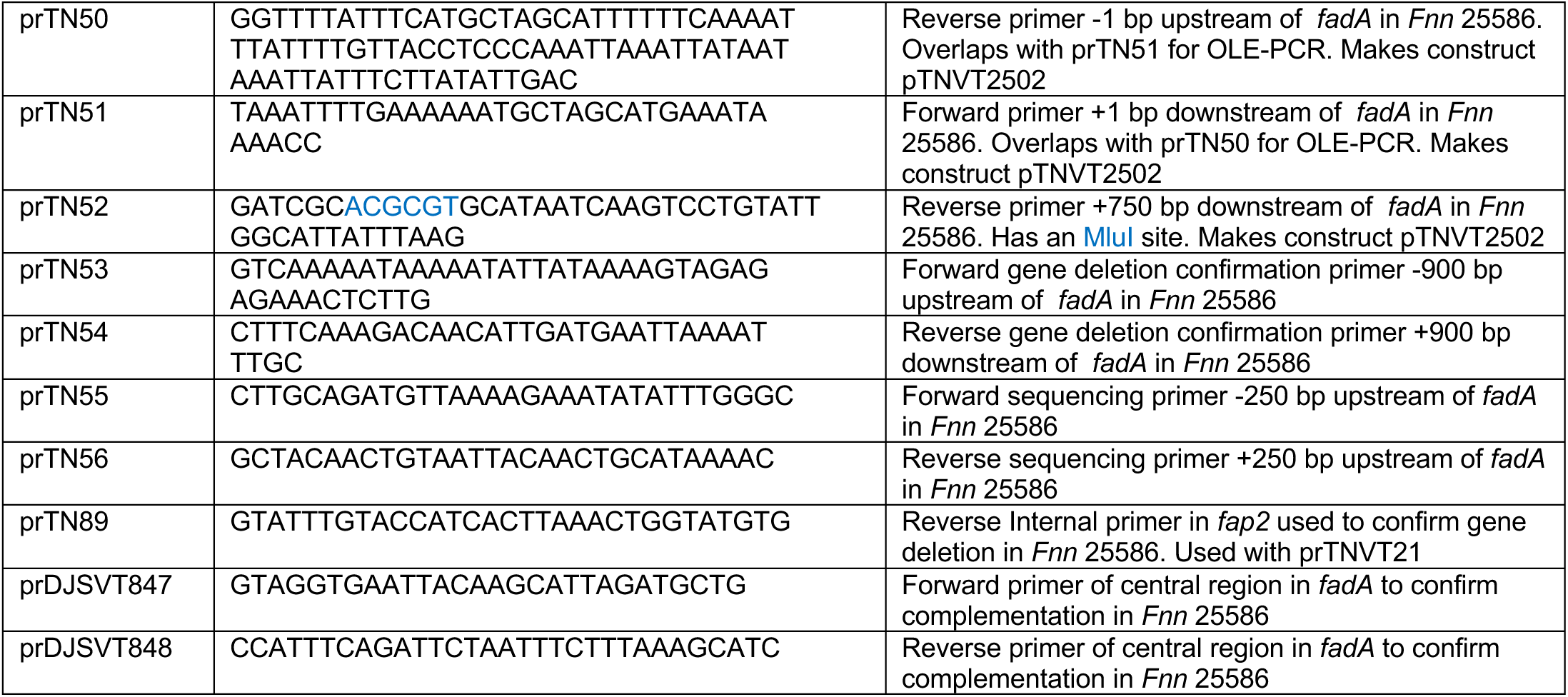
DNA oligonucleotides (primers) used in this study.

## Notes

### Competing Interest Statement

The authors have declared no competing interest.

